# Accounting for sensitivity of latent learning to behavioral statistics with successor representations

**DOI:** 10.1101/2025.03.01.640768

**Authors:** Matheus Menezes, Xiangshuai Zeng, Sen Cheng

## Abstract

Latent learning experiments were critical in shaping Tolman’s cognitive map theory. In a spatial navigation task, latent learning means that animals acquire knowledge of their environment through exploration, such that pre-exposed animals learn faster on a subsequent learning task than naive ones. This enhancement has been shown to depend on the design of the pre-exposure phase. Here, we hypothesize that the deep successor representation (DSR), a recent computational model for cognitive map formation, can account for the modulation of latent learning because it is sensitive to the statistics of behavior during exploration. In our model, exploration aligned with the future reward location significantly improves reward learning compared to random, misdirected, or no exploration, as reported by experiments. This effect generalizes across different action selection strategies. We show that these performance differences follow from the spatial information encoded in the structure of the DSR acquired in the pre-exposure phase. In summary, this study sheds light on the mechanisms underlying latent learning and how such learning shapes cognitive maps, impacting their effectiveness in goal-directed spatial tasks.

**Author summary:** Latent learning enables animals to construct cognitive maps of their environment without direct reinforcement. This process facilitates efficient navigation when rewards are introduced later, that is, animals familiar with a maze through prior exposure learn rewarded tasks faster than those without pre-exposure. Evidence suggests that the design of the pre-exposure phase significantly impacts the effectiveness of latent learning. Targeted pre-exposure focused on future reward locations enhances learning more than generic pre-exposure. However, the underlying mechanisms driving these differences remain understudied. This study investigates how pre-exposure methods influence subsequent navigation task performance using an artificial agent based on deep successor representations — a model for learning cognitive maps — within a reinforcement learning framework. Our findings reveal that before reward learning, agents receiving targeted pre-exposure develop spatial features more closely aligned with those of agents learning from rewards, compared to agents experiencing generic pre-exposure. This alignment enables the targeted pre-exposure agent to take more effective targeted actions, resulting in accelerated initial learning. The persistence of this advantage, even when modifying the agent’s exploration policy, indicates a robust cognitive map within the successor representation.

## Introduction

Latent learning demonstrates that animals can learn information about the spatial layout of their environment without forming specific associations between spatial locations and rewards and that this spatial information speeds up learning when rewards are introduced later [1, 2]. These types of experiments have played a crucial role in shaping Tolman’s cognitive map theory [3]. Notably, environmental constraints do not prevent latent learning from occurring. For example, Tolman incorporated doors into his latent learning experiments [3] and Daub tested their effect — the inability of animals to retrace their steps and only move forward — on the maze’s learning criteria [4]. Latent learning was observed no matter the structure of the environment.

Nevertheless, the design of the pre-exposure has a significant impact on reward learning of rats in the Dashiell maze [5]. Karn and Munn compared the learning performance of five groups of animals. The first group received no pre-exposure and behavior was reinforced immediately when animals were introduced to the maze (direct learning). In their study, the second and fifth groups’ pre-exposures focused on habituation to the confinement without explicit maze exploration. The third group explored the maze for five consecutive days, with trials ending when the animal reached the empty place where the reward would be provided later in the following experimental phase (targeted pre-exposure). The animals in the fourth group also explored once a day for a duration matched to that of the third group and were removed from any location in the maze (continuous pre-exposure). It was found that targeted pre-exposure facilitated learning and improved animals’ performance significantly more than continuous pre-exposure did. The continuous pre-exposure group in turn learned more effectively than the direct learning and the groups without explicit exploration.

Consistent with the observations by Karn and Munn, a study by Sutherland and Linggard found a similar pattern in rats that were trained to swim to an invisible platform in the Morris water maze [6]. The animals were divided into three groups: one was repeatedly exposed to the correct platform location (targeted pre-exposure), another to an incorrect location (which we name “mistargeted pre-exposure”), and the third to an unrelated water basin in a different room (naive group, direct learning). Before being exposed to the platforms, the targeted and mistargeted pre-exposure groups had a 30-second window for swimming around the water maze, while the naive group was allowed to swim around the unrelated water basin. During the (rewarded) learning phase, animals were placed on an arbitrary starting platform, followed by 90 seconds to complete the task. The targeted pre-exposure group rapidly acquired the spatial location of the escape platform, easily surpassing the performance of the other two groups. Intriguingly, even though the mistargeted pre-exposure group took longer to locate the platform than the naive group initially, they quickly outperformed the naive animals after learning the correct location.

To account for the above experiments, it has been hypothesized that animals built a cognitive map during the exploration in the pre-exposure phase. The cognitive map is thought to encode spatial relationships within the environment [3, 7, 8], which enhances learning to navigate to the reward location. Guo and colleagues have suggested that the effectiveness of the encoding occurs after repeated exposures to the environment and sleep in-between [9]. They found that hippocampal cells in mice did not quickly develop into strongly spatially tuned place cells, but remained weakly tuned to space and gradually developed correlated activity with the other neurons. They propose that weakly spatially tuned cells build connections that link discrete place fields to a map. In another recent study, Scleidorovich and colleagues proposed a computational framework for latent learning based on hippocampal replay [10]. They showed that targeted pre-exposure improves the agent’s learning phase when updating a state-action path matrix such that it reinforces the connectivity toward the goal location. However, this agent failed to show latent learning during continuous pre-exposure. In summary, the influence of diverse experimental designs on the outcomes of latent learning and the development of the cognitive map remain poorly understood.

Reinforcement learning (RL), specifically a version of the successor representation (SR) algorithm [11] supported by deep neural network and memory replay has been used to model latent learning [12]. The SR learns a spatial relationship between states in an environment independent of a reward function, allowing learning in an environment when no rewards are provided. The SR has consistently supported behavioral and neural findings [13–15] and outperformed other RL algorithms, such as model-free and model-based methods [11, 15, 16]. While it is well-known that the SR is sensitive to the agent’s policy [15], i.e., the statistics of the agent’s behavior, the impact of this sensitivity on modeling behavior and learned spatial representations have received little attention.

Here, we study whether and how this property of the SR accounts for differences in later learning between targeted, continuous, mistargeted, and no pre-exposure. Our simulations reveal several insights. First, each pre-exposure design indeed shapes a distinct SR, which in turn drives differences in behavior. Second, restricting spatial behavior, such as by inserting doors that open in only one direction, facilitates SR acquisition during pre-exposure, leading to faster escape latencies in subsequent learning, especially in time-constrained tasks, such as ones with a restricted small number of allowed steps. Finally, the SR’s influence on latent learning is robust across diverse state representations (one-hot vs image-based) and RL exploration policies (softmax vs *ϵ*-greedy), underscoring the SR’s enduring impact regardless of the specific exploration strategy animals might employ.

Overall, we take a step forward in understanding how the SR can not only manifest latent learning, but also how the policy dependence of the SR accounts for sensitivity of latent learning to pre-exposure design and movement restrictions. This contributes to a deeper understanding of how animals’ learning of cognitive maps is critically modulated by their spatial behavior.

## Materials and methods

### The successor representation in reinforcement learning

We study latent learning in the framework of RL where an agent interacts with an environment by taking an action *a* to transition from state *s* of the environment to another state *s′* with the probability *p*(*s′* |*s, a*). The agent chooses actions according to a policy *π*(*a*|*s*) and receives an instant reward in each state *R*(*s*). The goal of the agent is to maximize the total amount of reward. One way to achieve this is based on the state-action value function, also called the *Q* function, which is formally defined as the expected cumulative discounted reward under the policy *π*:

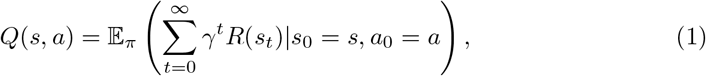

where *γ* is the discount factor. During decision-making, the action with the highest *Q* value will be selected with the highest probability. To maximize the total reward, the agent has to find the optimal function *Q*^∗^, which has to satisfy the Bellman equation [17]:

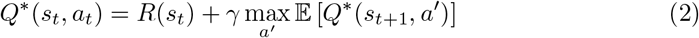

There are two major classes of RL algorithms for solving this problem: model-free and model-based. Model-free approaches directly learn an action policy on the data collected from the interactions with the environment. They usually leads to a good policy, but are data-hungry and require many interactions with the environment. They also struggle to generalize to new tasks and need to relearn the policy when the environment dynamics change. By contrast, model-based approaches learn a model of the dynamics of the environment and leverage the model to optimize the policy. This makes them more data-efficient, but increases the complexity of the computation and is prone to accumulate errors due to inaccuracy of the model, particularly with large state spaces. A third approach that lies on a middle ground between data-efficiency and model complexity is the successor representation (SR) [11].

The SR algorithm estimates the expected frequency of transiting from a certain state to successor states [14]. Specifically, one row of the transition matrix *M* represents the expected visitation frequencies from the current state to all states. *M* can be expanded by one extra dimension to incorporate actions, such that given a state *s*, an action *a*, and future state *s′, M* is defined as:

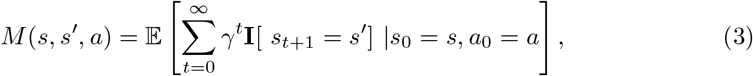

where **I**[.] = 1 when its argument is true and 0 otherwise. Note, *M* is independent of any reward function, enabling rapid adaptation to changes in the environment’s reward structure. In this sense, SR agents are more robust to changes than model-free methods, as the agent only needs to relearn the new reward function *R*(*s*) while maintaining the existing transition model *M*, resulting in fast integration of changing reward functions in dynamic environments.

The transition matrix has to satisfy a Bellman equation:

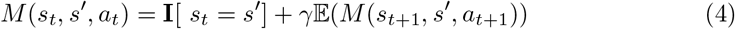

where *a*_*t*+1_ is selected based on the running policy. As the agent acquires new experiences over steps, *M* (*s, s′, a*) is updated with a temporal-difference learning rule [17]. Assuming the agent goes from *s*_*t*_ → *s*_*t*+1_by taking action *a*_*t*_, then the agent can implement the learning rules:

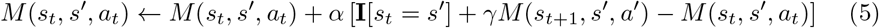

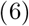

Given the SR, the *Q* value function of selecting action *a* at state *s* can be computed by summing the product of *M* (*s, s′, a*) and the immediate reward for each corresponding successor state [11]. This process captures the anticipated long-term value of the state-action pair based on SR’s predictions about future interactions [13]:

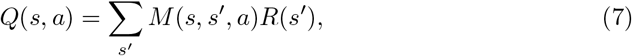

### Decision-making with deep successor representation

In complex environments, e.g., with high-dimensional states and continuous actions, tabular methods become infeasible and deep neural networks are used to approximate value functions [18]. Kulkarni and colleagues [19] combined deep learning with SR algorithm by learning the spatial and temporal relationship between states with deep neural networks, and representing the reward function as a trainable weight vector. We refer to this agent as the deep successor representation (DSR) in the rest of this paper.

The key concept in DSR is the successor feature (SF), denoted as *m*_Φ_(*s, a*) where Φ refers to the parameters of the neural network, which represents the expected future occupancy of the state features, starting from the state-action pair (*s, a*):

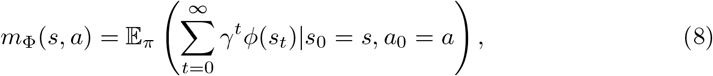

where *ϕ* is a feature extraction function. When the reward function is a linear function of the state feature: *R*(*s*) ≈ *ϕ*(*s*) · **w**, the *Q* function can be decomposed as follows:

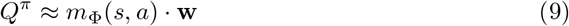

with **w** being a weight vector. We implement **w** using a one-layer, linear neural network with no biases, whose input is the state feature *ϕ*(*s*).

Throughout our simulations, we implement a version of DSR as the running algorithm to model the effects of latent learning [12, 20]. We employed a simple replay mechanism where all visited (state, action, next state, reward) tuples are equally sampled from the memory buffer and replayed to update the neural networks. Additionally, for the non-continuous designs, we explicitly replayed transitions from the goal state to itself. This procedure simulates the period described in experimental studies, during which animals are confined in a goal cage for a short period, thereby manipulating the likelihood of associating the goal with rewards.

The agent learns a separate SF network for each action, whose input is a state feature and output a SF vector, *m*_Φ_(*s, a*), which captures the future visitation to all states, given the agent is in state *s* and chooses action *a*. Based on Eq. 8, each SF vector can be approximated as a weighted sum of state features:

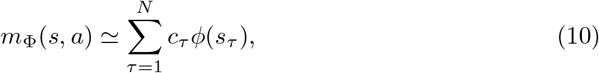

where *N* is the number of states and the coefficients *c*_*τ*_ represent the expected future visitation of state *s*_*τ*_ from the state-action pair (*s, a*). For each action, we take the SF vectors from all states and turn the above equation into a matrix format:

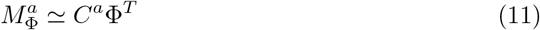

Hence, a row *i* in *C*^*a*^ represents the expected visitation frequencies from the current state *s*_*i*_ to all states. The matrix *C*^*a*^ essentially represents the same information as the matrix *M* in in tabular SR (Eq. 3). We can solve the above equation as follows:

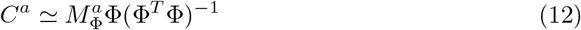

The accuracy of the solution depends on the matrix inversion (Φ^*T*^ Φ)^−1^. Ideally, the state matrix Φ should be full-rank, which means the state features should be linearly independent. For one-hot encoded states, *C*^*a*^ can be perfectly solved. For image-based inputs, we tried to extract the features from images such that they have minimal correlations. With *C*^*a*^, we can map both one-hot and image-based representations to the environment’s topology, with an entry *i* in each row corresponding to the expected visitation frequency of state *i* from the current state. Throughout this paper, we refer *C*^*a*^ as the deep SR matrix.

### Latent learning simulation setup

In T-maze experiments by Blodgett and Tolman [1–3], rats exposed to the maze without explicit reinforcement, such as food, initially exhibited no apparent decrease in error (entering dead alleys) and improvement in performance in reaching the goal box. However, when food was introduced at the goal location from 3 to 11 days later, they displayed a significant advantage over those who had not explored the maze and had received food at the goal from the beginning of the experiment.

All simulations were conducted in virtual environments designed in the CoBeL-RL (Closed-loop Simulator of Complex Behavior and Learning based on Reinforcement Learning and deep neural networks) framework [12]. CoBeL-RL allows the creation of a VR environment for trial-based experimental designs, where each trial consists of agent-environment interactions referred to as steps. Each step in CoBeL-RL corresponds to an experience tuple (*s, a, r, s′*), which the agent can either use for online learning or store in a memory structure for later replay.

We built two environments to simulate latent learning: a 10 × 10 open grid-world (Fig. 1a left) and a 14-unit T-maze (Fig. 1a right) similar to those used by Tolman. Each environment has either a one-hot encoding or an image-based state representation. Each environment had a fixed goal (g, green color) and starting (s, red color) state, with the Tolman maze having 72 or 30 states for one-hot encoding or image-based representation, respectively. We extract the state’s features encoding the images using the VGG16 network [21] (Fig. 1b), with an RGB image input of (32 × 128 × 3) and an output of (1 × 4 × 512) units. This representation makes it is possible to distinguish each feature (Fig. 1c) and compute its contribution to the successor features of other states by Eq. 12.

**Fig 1.**
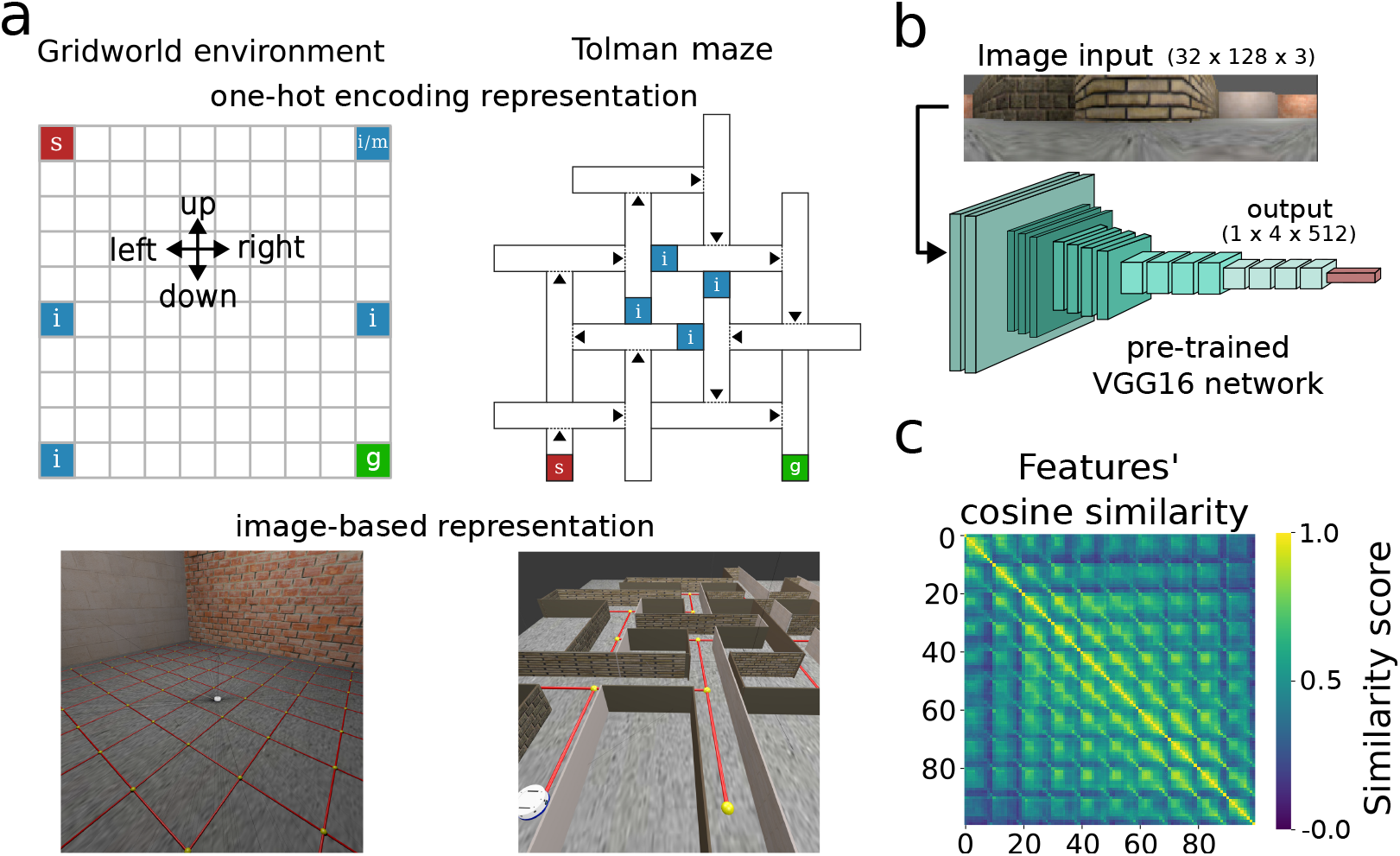
Simulation environments. **a**) Spatial setup of the environments. The gridworld is a 10 × 10 arena for both one-hot encoding (top-left) and image-based (bottom-left) representations. The Tolman maze [3] is either a 72-state environment (top-right) with one-hot encoding representations, or a 30-state environment with image representations (bottom-right). In the Tolman maze, virtual doors (dotted lines), if activated, can restrict the agent’s movement to only one direction (along arrow). The starting and goal states are indicated by s and g, respectively. The states marked with i are the additional starting locations during the continuous pre-exposure, while m is the incorrect goal location in the mistargeted pre-exposure. The locations of these distinct states shown in the one-hot encoding setting are the same in the image-based environments. **b)**. In the image-based learning, a pre-trained VGG16 network [21] encodes a (32 × 128 × 3) RGB image into a (1 × 4 × 512) state feature that is further used as the input to the deep SR networks. **c)** Cosine similarity between each pair of state features in the gridworld. Features of closeby locations have high similarities, but diverge for distant locations. In this, and comparable plots, the 10 × 10 states are stacked to a 100-dimensional vector to facilitate the plotting of similarity between all states.

To navigate, the agent selected one of four actions (up, right, down, or left). We adopted an action mask that disallowed the section of actions that would violate the geometric constraints of the environment. This prevented the agent from getting stuck in the same state, thus leading to a more dynamic exploration. Furthermore, we incorporated virtual doors (dotted lines) in the Tolman maze with the one-hot encoding representation such that, when enabled, they constrained the agent’s movement in one direction (along the arrow).

We tested our DSR agents in learning tasks with or without pre-exposure. In direct learning, just as in the rewarded control group in Tolman’s experiment, the agent receives a positive rewards (*R* = 5), if it reaches the goal location. Each trial begins at the start location. By contrast, in the three latent learning conditions, the agent goes through a pre-exposure phase for 50 trials that are not rewarded at the goal location (*R* = 0). This was followed by 50 trials, in which the agent received a reward of (*R* = 5) at the goal location. Additionally, a maximum of 300 time steps was imposed per trial in the open gridworld, and 1500 or 150 in the Tolman maze for the one-hot encoding or image-based representations, respectively.

### Task designs

We set up three experimental designs for latent learning simulations. During targeted pre-exposure, trials terminated when the agent reached the unrewarded goal state. This is similar to the experiments coined as Blodgett type [7], and conducted by Tolman and other researchers [2, 3]. During continuous pre-exposure, the agent explored the empty maze for the maximum pre-defined time steps [5, 7], whether or not it reached the goal state. Moreover, the agent initiated a trial from multiple locations, including the goal state (Fig. 1, potential starting states are i,s, and g). The mistargeted pre-exposure studied only in the grid-world environments, and trials ended when the agent reached the “mistargeted” state m (the location that will be rewarded later in the experiment) [6]. The learning phase was identical for all groups: the agent started in state s, and a trial ended upon reaching either the goal g or the maximum number of steps, whichever occurred first.

To assess how direct and latent learning designs influence the agent, we monitor its variables using CoBeL-RL embed tools [12]. These include analyzing behavioral variables, such as escape latency and success rate, alongside the learned successor feature (SF) vectors and Q values. For the latter, we recorded the learned SF and Q values estimated by the agent’s network at different phases of training.

## Results

### Learning the spatial connectivity during exploration drives latent learning

We first compared targeted pre-exposure with direct learning. The agent selects actions based on an *ε*-greedy policy, with *ε* = 0.3. By the end of learning phase, both agents mastered the spatial task in the two environments and using the two types of representations (one-hot encoding and image-based), as shown by the error reduction. Notably, the latent learning agent outperformed the direct learning counterpart (Fig. 2a, b). These results qualitatively match the behavior of animals [3, 7] and previous computational simulations [10, 16], in which trajectory errors dropped as latent learning animals encountered rewards. Similarly, the escape latency of our latent learning agent notably decreased by the moment they experienced rewards (trial 0, blue curves, Fig. 2a). During pre-exposure (−50 to −1 trials), this agent exhibited notable variability across simulations (shaded area, escape latency), with a similar trend observed when measuring the agent’s success rate in reaching the goal state (Fig. 2b). The variability of performance during the pre-exposure is higher than during learning, indicating no discernible progress on the task during the pre-exposure phase. Nevertheless, once the reward was introduced, the success rate of the pre-exposed agent increased faster than that of the direct learning agent, an indication of latent learning.

**Fig 2.**
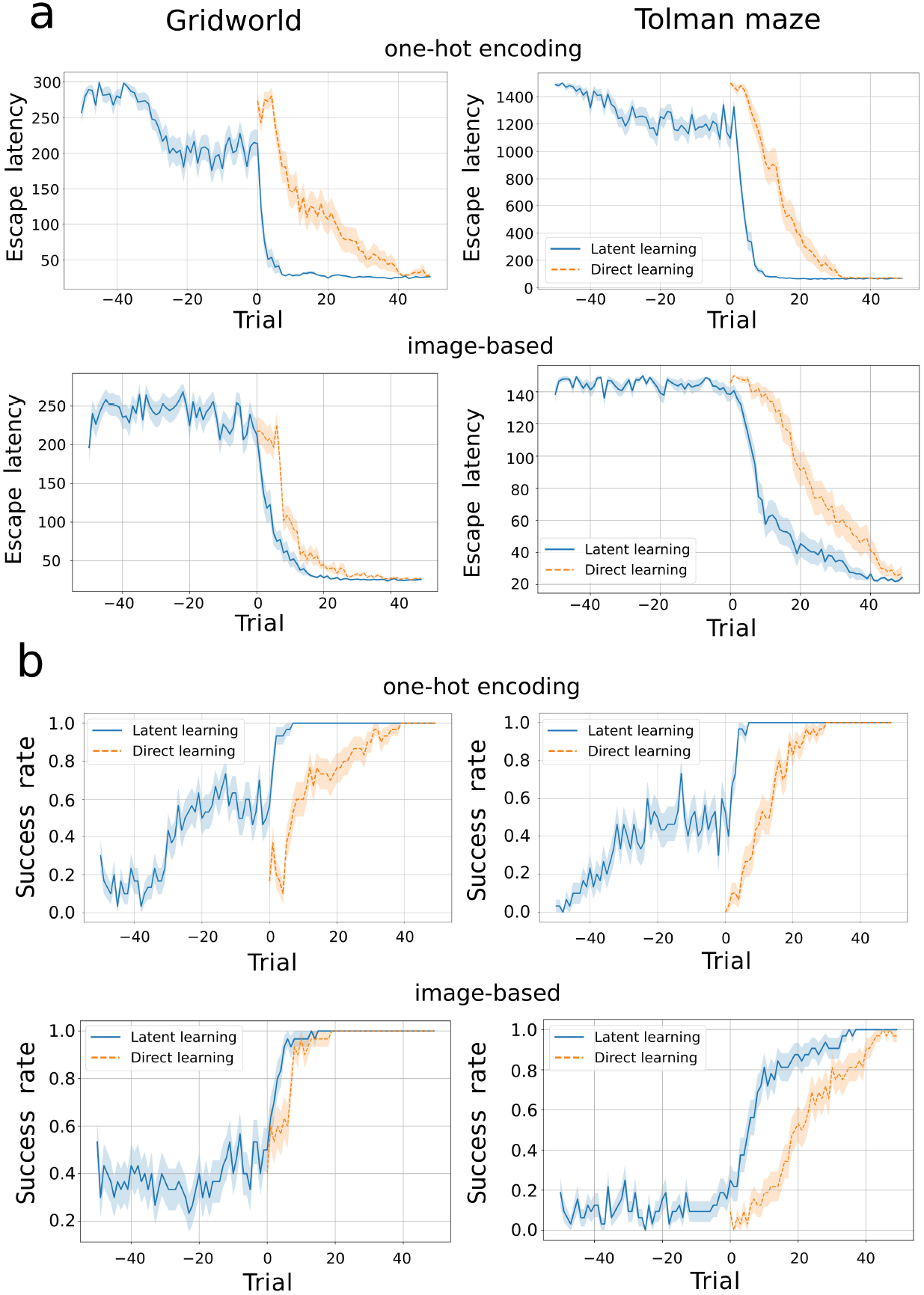
Learning curves demonstrating latent learning. Left column: gridworld. Right column: Tolman maze. **a**) Latent learning agents (blue curve) with targeted pre-exposure to the environment (trials the neural network. Although this50 to the neural network. Although this1) consistently reach the goal faster than direct learning agents (orange dashed curve), after a reward is introduced from trial 0 in both one-hot encoding (top row) and image-based representations (bottom row). Curves represent the averages over 30 simulations, and the shaded area the S.E.M. The success rate of reaching the goal location in each trial. Latent learning agents are mostly unsuccessful on the spatial task during the pre-exposure phase, but after the introduction of rewards they reach the goal state more consistently than the direct learning agents do.

The gradual drop in error rates in the pre-exposure phase might be due to the targeted pre-exposure design. Although receiving no rewards, the agent still slightly increased its expectation of the future occupancy of the goal location because the pre-exposure phase was terminated at the (future) goal state. With the learned reward function being near-zero for all states, but not exactly zero due to the nature of the minimization procedure, the tendency for the agent to go to the goal state was increased.

The deep SRs generated by the neural networks reveal the spatial structure learned by the agents in both one-hot (Fig. 3) and image-based representations (Supplementary Fig. 1). For the direct learning agent, the deep SRs at trial 0 reflect the random initialization of the network (Glorot uniform algorithm [22]). In contrast, for the latent learning agent the deep SRs at trial 0 show a pattern that captures the local connectivity of states with no directional bias. This was the result of a random exploration strategy during the pre-exposure phase. By the end of the learning phase, both agents have developed a similar deep SR structure that reflects a movement from the start to the goal location. This progression is further evident for the deep SR of individual states, where activity is inhibited to dead-end or backward states (Fig. 4, observed states are marked with orange crosses). Interestingly, the deep SR of individual states, as well as its matrix form, exhibits high values at the goal state after the learning phases (with noticeable patterns in the gridworld environment after pre-exposure), even for states distant from the goal. This effect arises from replay in our model, through which the SF is updated from the goal state to itself, amplifying its value and propagating it across other states through the neural network. Although this modeling choice is unconventional in RL, it proved necessary for learning in our simulations. Its broader implications — and potential links with experimental findings — merit further investigation.

**Fig 3.**
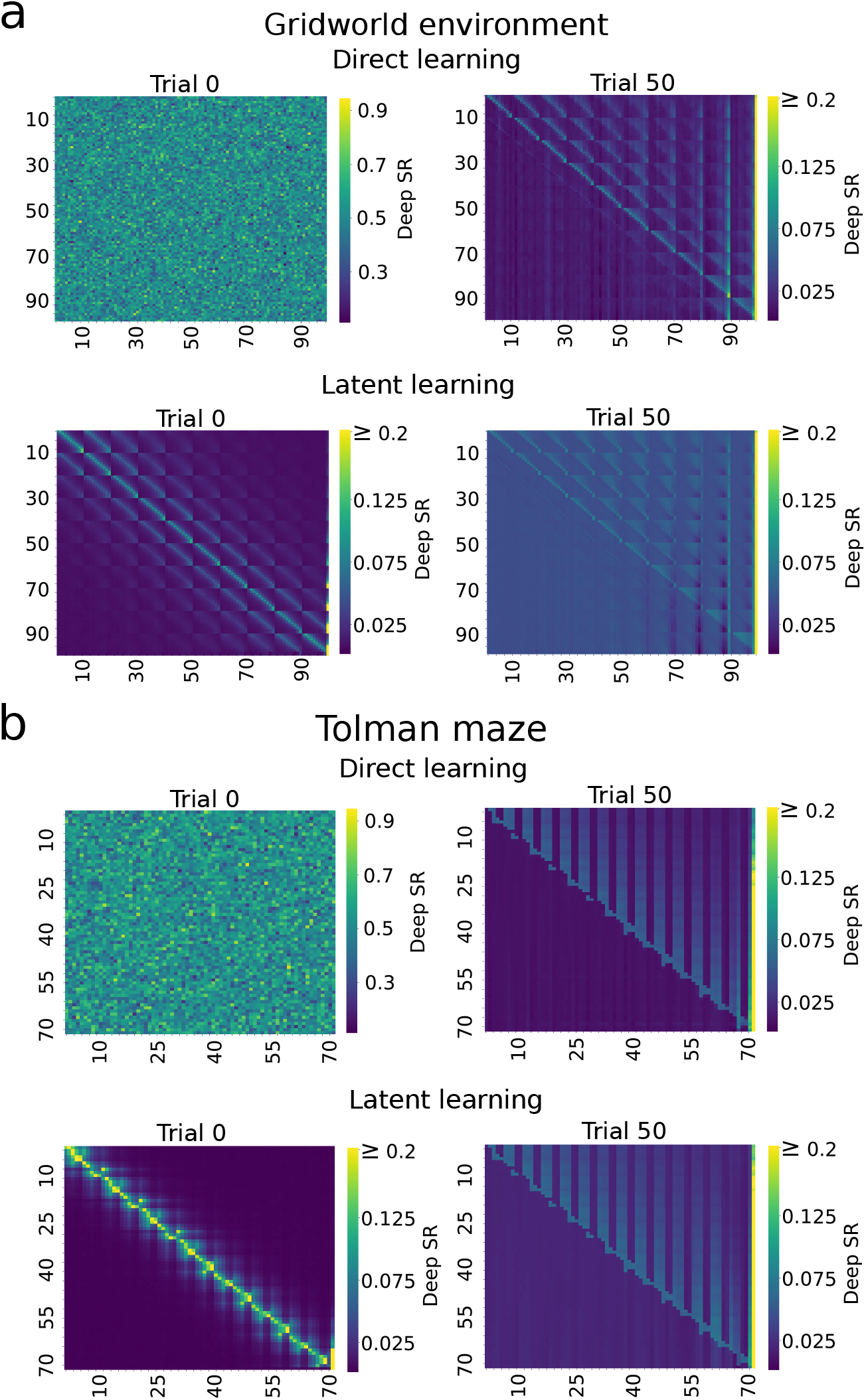
Agent learns local connectivity of the maze during latent learning. Deep SR matrices derived from one-hot encoding for all states are shown for the **a**) gridworld for the right action and **b**) Tolman maze for the up action, averaged over 30 simulations. Rows and columns correspond to the state indices. Each row of panels represents to the successor transitions of a particular state to every state in the environment. Trials 0 and 50 correspond to before and after the learning phase, respectively. At the end of the learning phase, both agents exhibit similar transition patterns that reflect a movement from the start to the goal location in both mazes.

**Fig 4.**
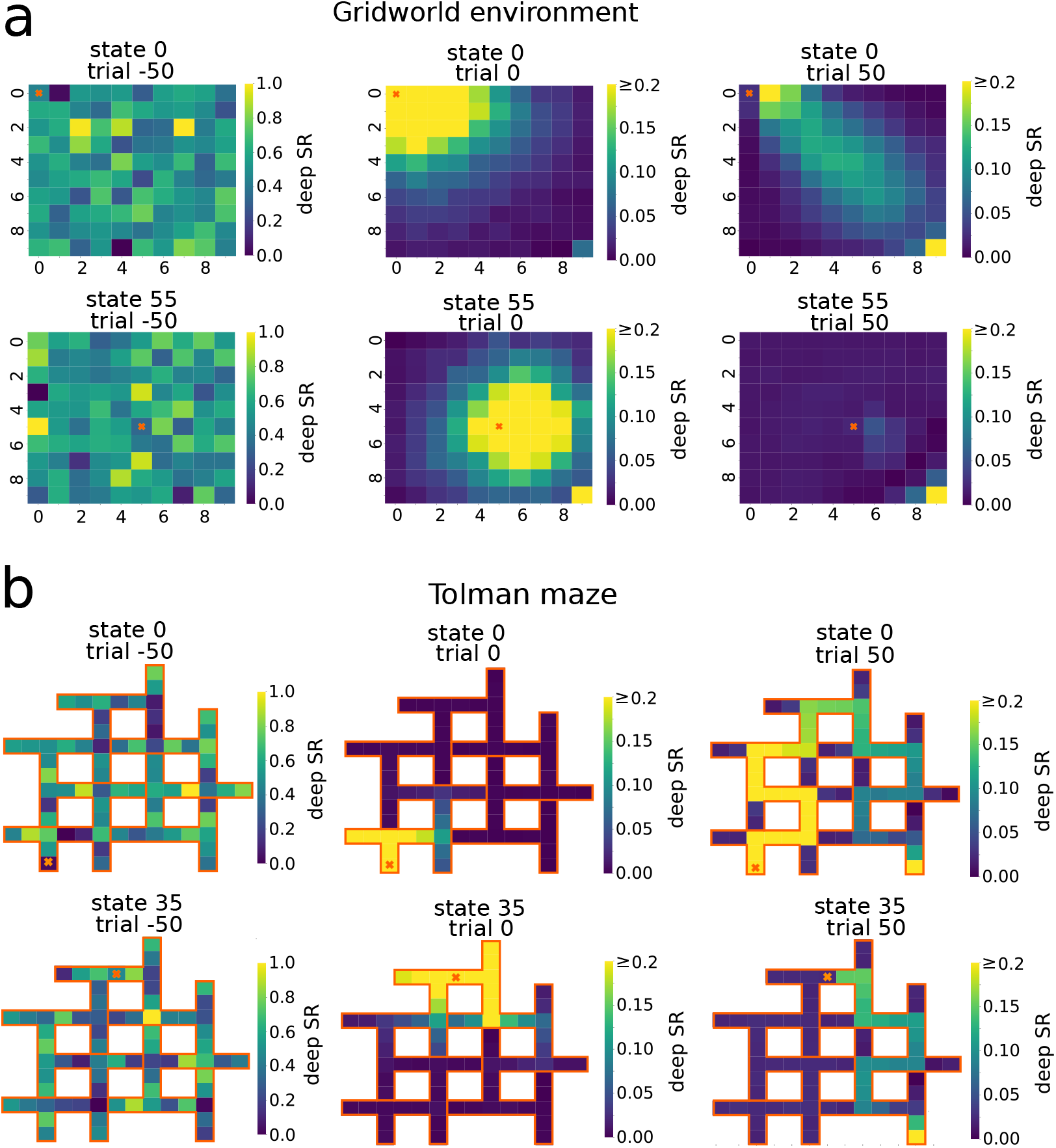
Successor representation of individual states reveal predictive encoding. Shown are visualizations of one row of the deep SR matrix (one-hot encoding, targeted pre-exposure), mapped onto the environment structure, for the state indicated by the orange cross in the **a**) gridworld and **b**) Tolman maze. Trial −50 corresponds to the randomly initialization. After the pre-exposure to the environment (trial 0), the future states which are expected to be visited by the agent are mostly near the current state, whereas by the end of learning (trial 50), the future occupancy of states has evolved into a path leading to the goal.

Using image representation to learn the deep SR matrix allows agents to generalize across new inputs that were not used during training (Fig. 5). This results in paths that direct towards the goal state when processing new inputs. In contrast, one-hot encoding representation does not exhibit a bias towards the goal for novel inputs. This happens because the one-hot encoding lacks relational structure across states, preventing the agent from inferring similarities between trained and unseen inputs. With this, our network outputs a representation that can be random or linked to a representation of a particular state, e.g. the starting location in Fig. 5, top row.

**Fig 5.**
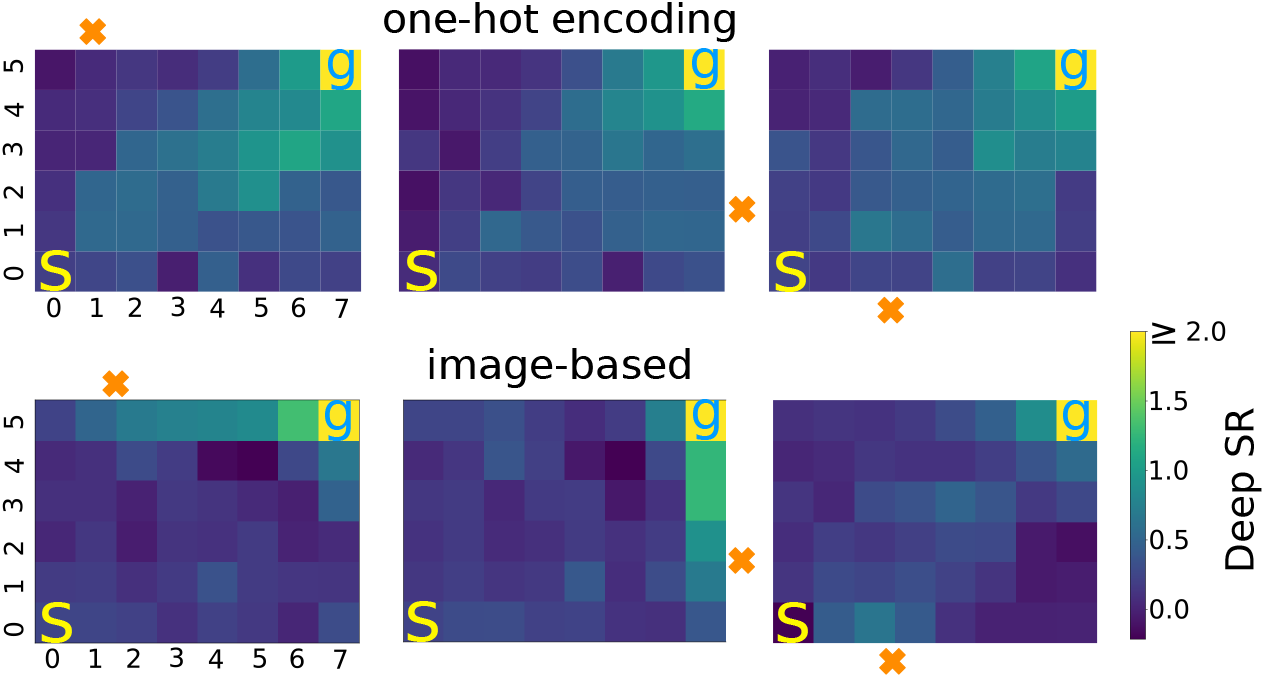
Agent generalizes only image-based state representation to novel inputs. Bottom: Deep SR matrix of individual states trained with image features and mapped into an 8 × 6 environment (direct learning, 20 simulations) shown for a location that was not used during training (orange x). Note, the clear representation of a trajectory toward the goal. s and g mark the start and goal states, respectively, that were used during training. Top: When one-hot encodings are used for state representations, the deep SR matrix exhibits no directional bias for the novel location toward the goal location.

Additionally, we examine the Q-values resulting from multiplying the successor features with the reward weight vector for the one-hot encoding representation (Eq. 7). The Q-values are initially stochastic for the direct learning agent due to its randomly initialized SF and the reward weight (Fig. 6, trial 0). This contrasts with the latent learning agent, which has acquired both the SF local connectivity and an approximate reward function (*R*(*s*) ≈ 0, for all *s*) after exploring the environment devoid of reward (trial 0 of latent learning). This leads to a more uniform distribution of Q-values across various regions of the environment. The color intensity of each arrow in the graph represents the normalized values (0 – 1) associated with each state-action pair. At the end of the learning trials (trial 50), the Q-values for both agents show prioritized actions that lead to the goal.

**Fig 6.**
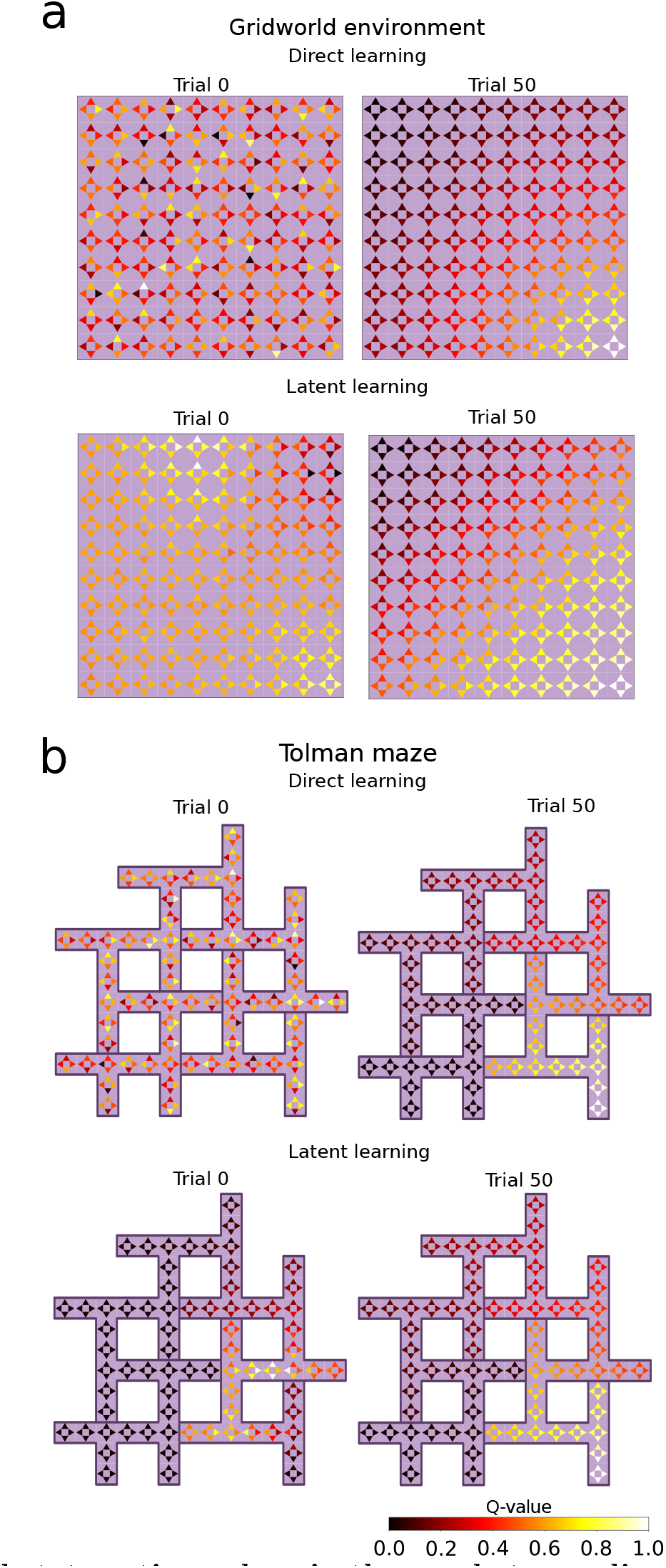
Learned state-action values in the one-hot encoding setup. For the **a**) gridworld environment and **b**) Tolman maze, the color of each arrow represents the Q value (indicated by color) of the corresponding action in that state. For direct learning agents, both the SF network and the reward weight vector are randomly initialized, resulting in a stochastic distribution of Q-values at trial 0. Latent learning agents have encoded the transition structure between neighboring states and a proper reward function (*R*(*s*) ≈ 0, for all *s*) by the end of the pre-exposure phase. This leads to a more uniform distribution of Q values, which nevertheless are sensitive to random fluctuations, e.g., there are high Q values near the starting in the gridworld where they should not be high. Following learning with reward, both agents converge towards similar Q values, with higher values for actions leading toward the goal.

### Impact of differences in task design on latent learning

Next, we investigated two different latent learning experimental design: continuous and mistargeted pre-exposure. These agents also exhibited latent learning, consistently outperforming the direct learning agent in escape latency (Fig. 7a, b). However, they initially required more trials to learn the task than the agent receiving targeted pre-exposure. In the learning phase, the mistargeted design shows inferior performance compared to all other designs in the initial trials (0 and 2) for image-based representation. These subtle differences in performance align with findings from the literature [5, 6], indicating that targeted pre-exposure may confer an advantage in terms of learning speed of latent learning.

**Fig 7.**
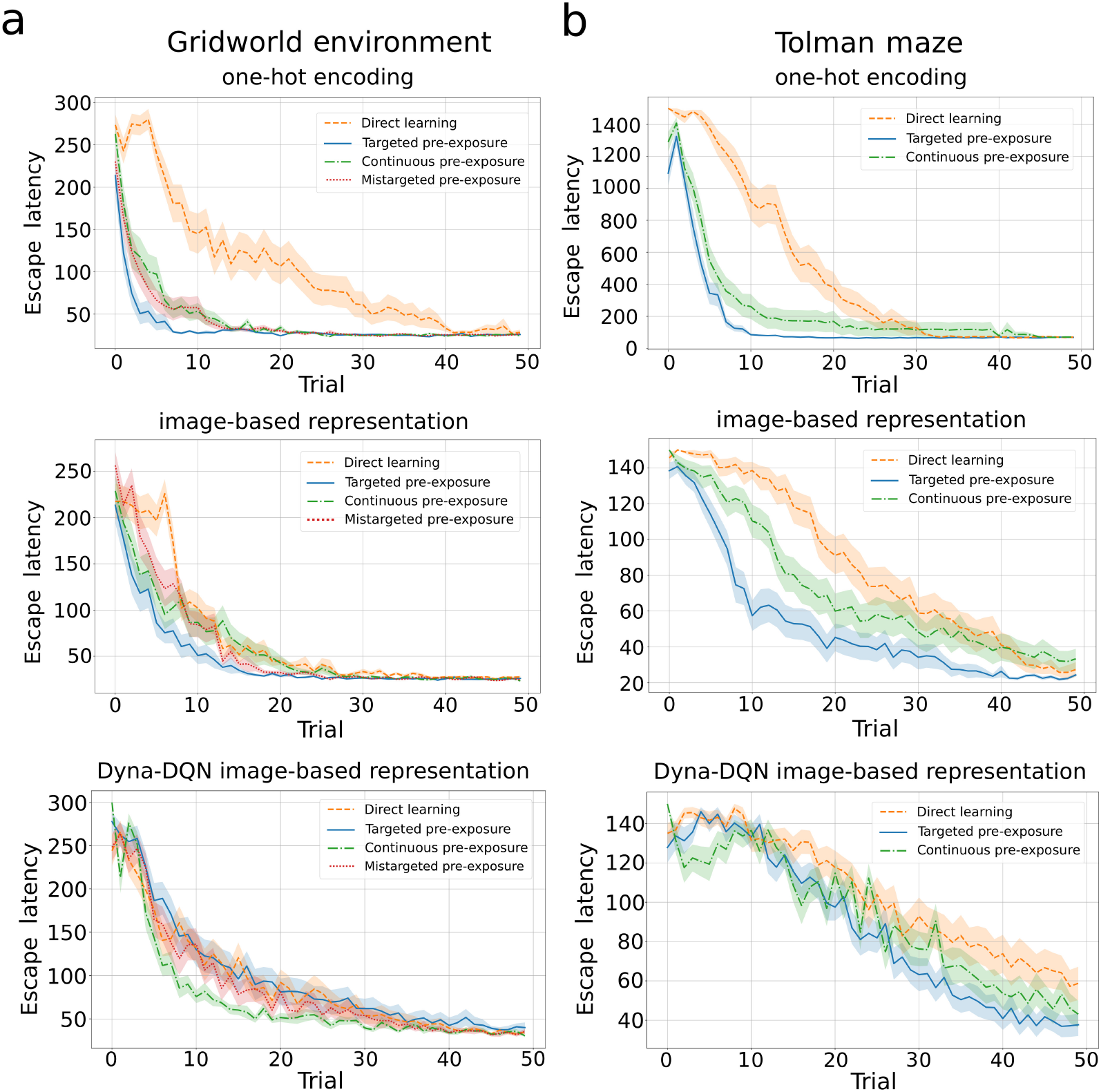
Experimental task design influences latent learning. Comparison of latent learning agents with three pre-exposure designs (targeted, continuous, mistargeted) to the direct learning agent for the gridworld. Lines indicate averages across 30 simulations and shaded regions the S.E.M. **a**) Performance in the grid world for agents using deep SR based on one-hot encodings (first row), deep SR based on image-representations (second row), and model-free image-based Dyna-DQN (third row). **b**) Same as in a) for the Tolman maze.

To examine the cause of the difference between the different pre-exposure paradigms, we compared the resulting SFs at the end of the exploration. Targeted pre-exposure led the agent to develop a spatial structure with higher similarity to the SF of the direct learning agent by the end of the rewarded learning than continuous pre-exposure did (Fig. 8). This higher similarity for the targeted pre-exposure agent was particularly pronounced for states closer to the goal, as estimated by the cosine similarity between the outputs of the SF network at each state for the direct learning agent and the latent learning agents (Fig. 8c,d). These findings indicate that even without rewards, the targeted pre-exposure helps the agent build spatial knowledge of the environment that favors movement towards the goal location, which potentially explains the observed faster learning compared to the continuous pre-exposure agent.

**Fig 8.**
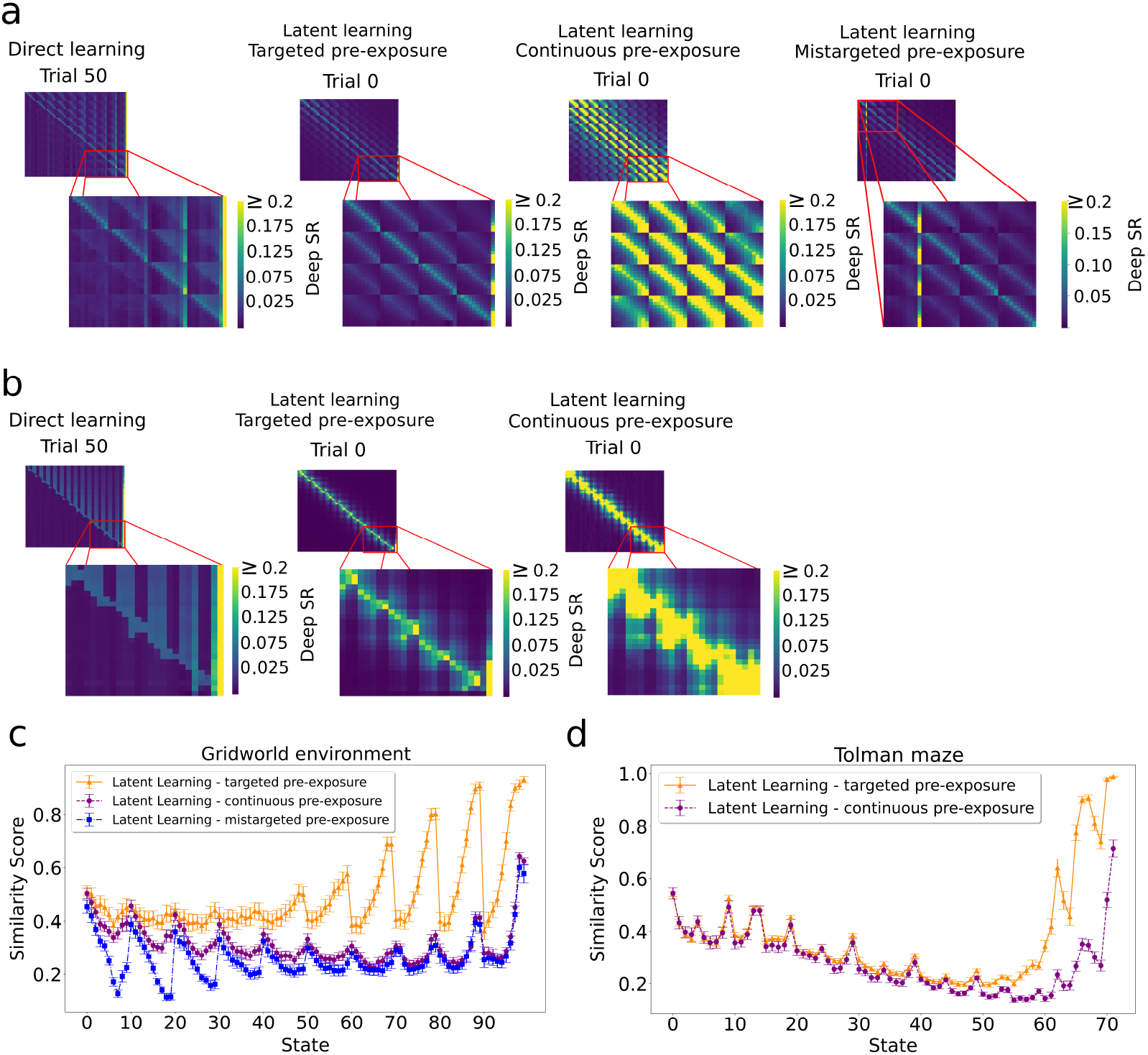
The structure of Deep SR accounts for performance difference in the three different pre-exposure paradigms. **a**) Left: Well-learned SF for the direct learning agent in the gridworld. Left-center: After targeted pre-exposure, the SF structure is similar to what the direct learning agent learns from rewards, particularly in states near the goal (shown in inset). Right-center: After continuous pre-exposure, the structure is more dissimilar to that of the direct learning agent. Right: After mistargeted pre-exposure, the structure is similar to that of the targeted pre-exposure, but for the mistargeted location. **b**). Same as in a) for the Tolman maze without the mistargeted pre-exposure simulations. **c**) For the gridworld. The cosine similarity between the SF vector from the latent learning agents at the end of the pre-exposure phase and that from the direct learning agent at the task’s conclusion. Targeted pre-exposure results in superior scores in states close to the goal (last state), while mistargeted pre-exposure shows the lower scores in states close to the mistargeted state. Error bars are S.E.M. **d**) Same as in c) for the Tolman maze.

To confirm this explanation, we further studied the RL policy that emerges from the SF just after the pre-exposure phase. To this end, we computed the Q values (Fig. 9a,b) by combining the agents’ SF and an approximate ground truth reward function *R*(**g**) ≈ 1, where **g** represents the goal state, and *R*(**s**) = 0 for all other states. In contrast to continuous pre-exposure, these values are higher for the targeted pre-exposure agent in states further from the goal — which facilitates the agent selecting the correct action when the learning phase begins. Furthermore, we compared the optimal actions for the task in the Tolman maze with the actions selected by the latent learning policies in the last trial before introducing the reward (Fig. 9c, trial 0). The learned policy selects the actions that lead to the goal state more frequently in most states for the targeted pre-exposure agent. In addition, the optimality ratio reveals that targeted pre-exposure consistently selects more optimal actions than continuous pre-exposure over all trials (Fig. 10). Our results suggest a pre-existing bias that carries over to the learning phase.

**Fig 9.**
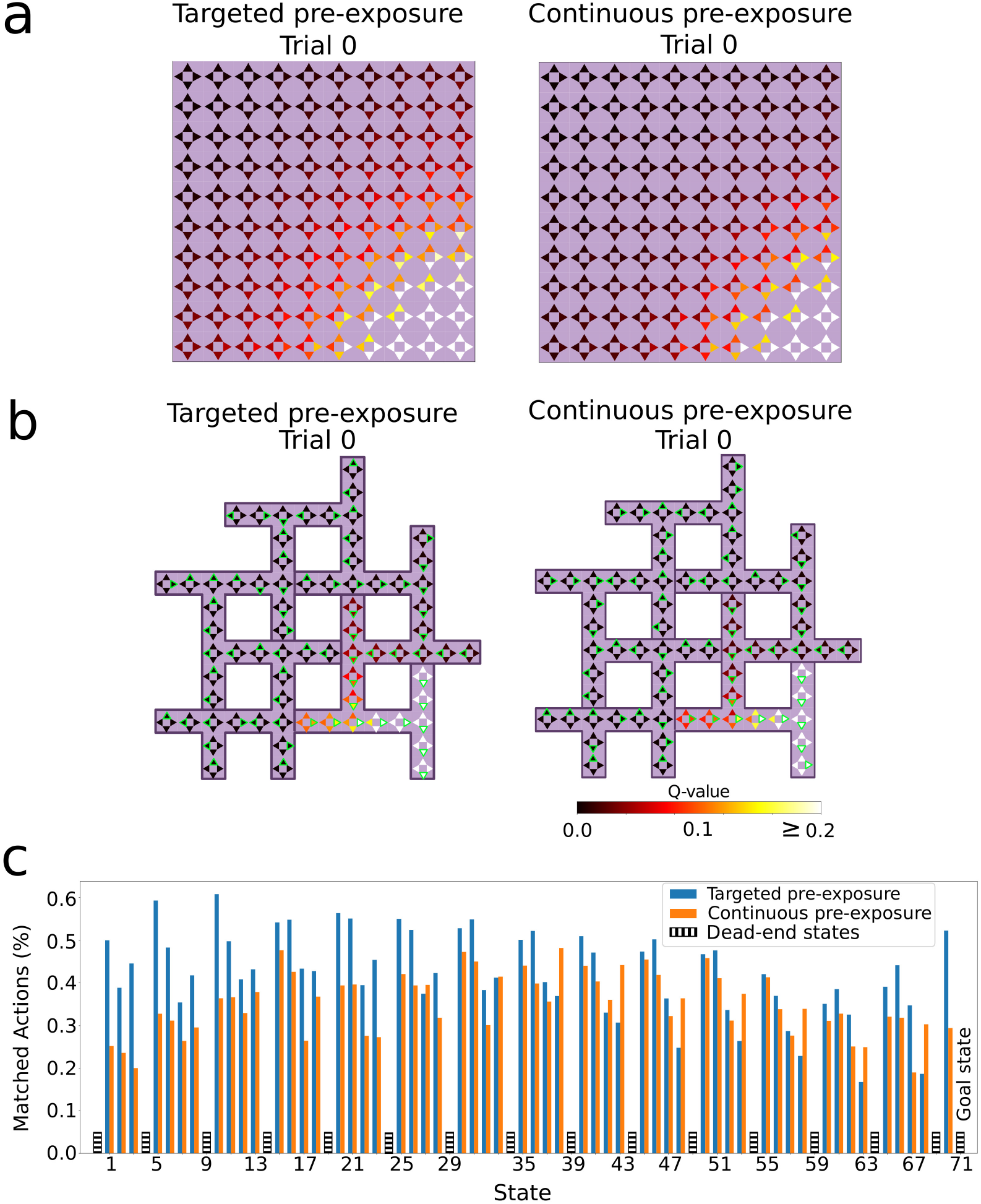
Evaluation of the policies learned during the pre-exposure phase. Q values for targeted and continuous pre-exposure in the **a**) gridworld environment, and **b**) Tolman maze, computed from the learned SF multiplied with the ground-truth reward function. Targeted pre-exposure drives elevated values for states that are more distant from the goal, compared to continuous pre-exposure. The green edges show the action with the highest Q-value in that state. **c**) The probability of selecting the optimal action at the conclusion of the pre-exposure phase in the Tolman maze, averaged over 30 simulations. Targeted pre-exposure tends to lead to the optimal actions more frequently than continuous pre-exposure. Dead-end states, where only one action is viable, exhibit an action selection probability of 1 (see Methods) and are omitted here for clarity.

**Fig 10.**
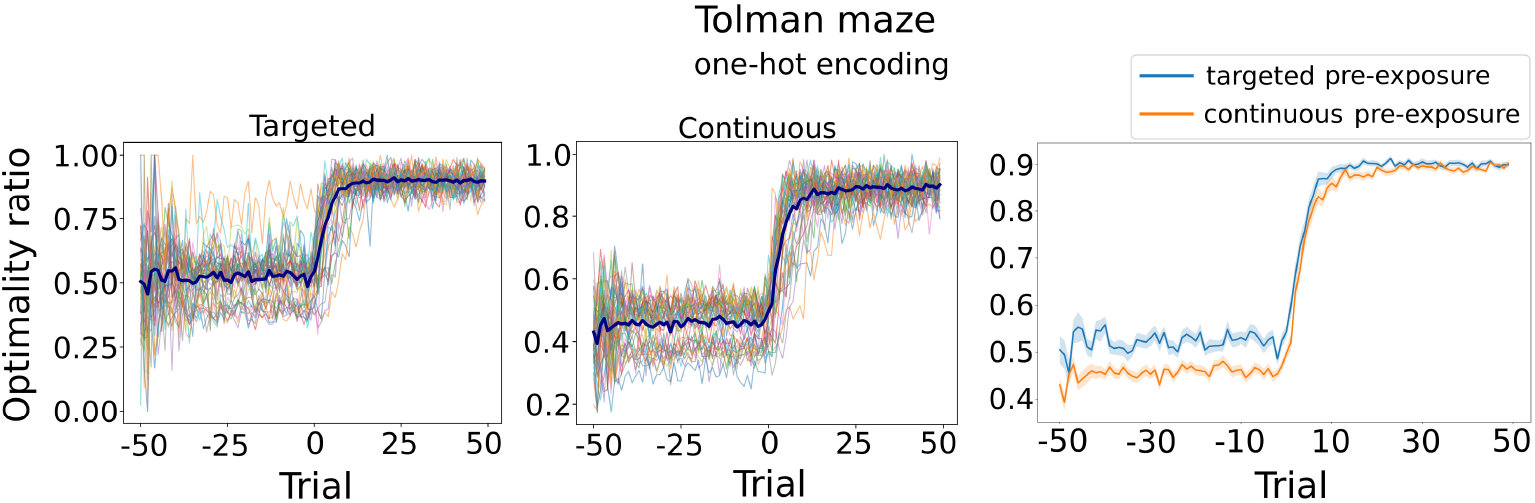
Targeted pre-exposure drives more optimal action selection than continuous pre-exposure in Tolman maze. Left and middle: Thin lines represent the optimality trace of a single simulation. For each trial, optimality was quantified as the ratio of optimal actions to the total number of actions taken across all states that transition to the goal, i.e., excluding dead-end states. Thick lines represent the average over 30 simulations. Right: The agents’ optimality ratios show that targeted pre-exposure consistently outperforms continuous pre-exposure in all trials. Average over 30 simulations, shaded regions indicate S.E.M.

To demonstrate that the sensitivity to the pre-exposure design indeed arises from the policy dependence of the SR, we implemented an RL agent based on the CoBeL-RL’s Dyna-DQN and model-based [23] using image-based representations. We used the same simulation parameters as for the deep SR agent. The performance differences that arise from pre-exposure designs in latent learning are not observed in model-free and model-based algorithms (Fig.7, third and fourth row). This is an indication that accounting for the different effects of pre-exposure is not a generic property of RL, but requires the specific properties of the SR.

### Effect of exploration strategies on latent learning

To investigate how robust the subtle learning speed differences are that emerge from different latent learning designs, we studied different RL exploration strategies (Fig. 11a). The targeted pre-exposure agent learned the task faster when using the *softmax* policy (similar to results with *ε*-greedy policy) using inverse-temperature *β* = 1, followed by continuous and mistargeted pre-exposures and direct learning agents. Nonetheless, the last three agents also significantly progressed in the learning phase, narrowing the performance gap observed under the *ε*-greedy policy (Fig. 8a,b).

**Fig 11.**
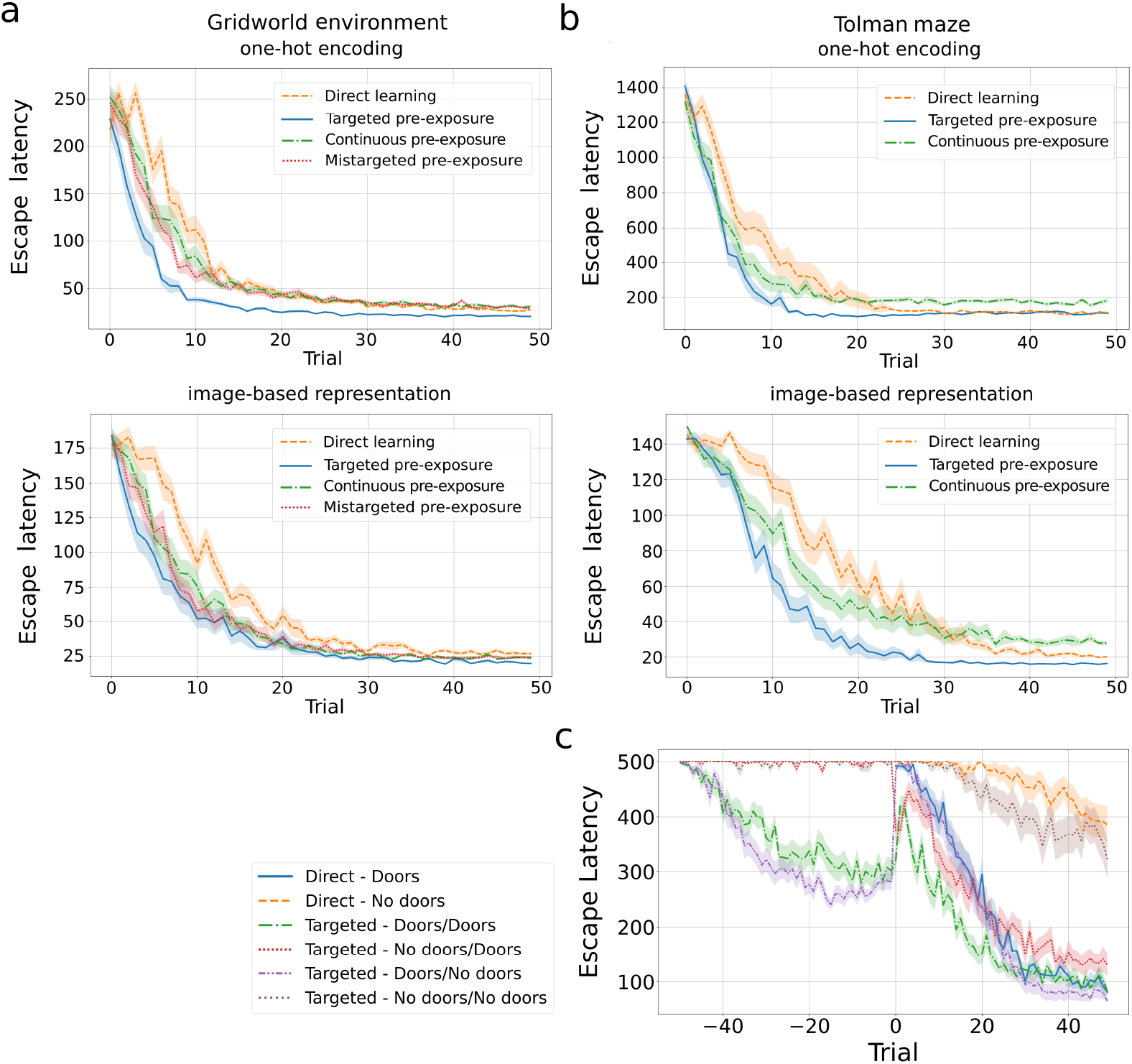
Latent learning observed with different RL exploration strategies. Evaluation of the softmax policy for exploration in the **a**) gridworld and **b**) Tolman maze (average of 30 simulations, error bars are S.E.M). Latent learning agents consistently exhibit faster learning during the learning phase, compared to direct learning agents. The subtle performance differences between targeted, continuous, and mistargeted pre-exposures persist even as the exploration policy changes. **c)** Restricting the agents movement, by introducing doors that are only passable in one direction, in the Tolman maze during pre-exposure and learning phases significantly influences the performance of the agent (average over 30 simulations, with a maximum of 500 steps per trial, error bars are S.E.M.). Performance is improved when doors are active. Legend labels for agents with pre-exposure indicate the door conditions during pre-exposure and learning phases, e.g. “No doors/Doors” indicates that doors were not active during pre-exposure, but were during learning.

We account for this performance gain under the softmax policy, as compared to *ε*-greedy policy, as follows. After exploration, Q values tend to become more uniformly distributed (see Fig. 9). If Q values of different actions in the same state are similar, the softmax policy chooses stochastically between these actions with almost equal probability, depending on the inverse temperature *β*. This leads the softmax policy to prioritize exploration in the initial states, potentially guiding the agent to the goal state faster during early learning. In contrast, the *ε*-greedy policy selects actions with the highest Q value with probability (1 − *ε*), even if the differences in Q values are small. This reduces exploration and might cause the policy to select the same action repeatedly during its greedy steps. This is especially detrimental if Q values for non-optimal actions leading to dead-ends are initially the highest.

Next, we examined the impact of physically constraining the movement of the agent on latent learning by using doors. The doors were set up in Blodgett’s and Tolman’s experiments [1, 3] to speed up learning by preventing the animals from revisiting mapped locations. Daub further studied the effects of doors by comparing latent learning in environments with and without doors [4] using four groups of animals in a 2 × 2 design. The groups were either pre-exposed to the environment without doors or not pre-exposed. In the learning phase, the door were either engaged or kept open. If no doors were used, animals that received pre-exposure (corresonding to “Targeted - No doors/No doors” in Fig. 11c) performed better than animals without pre-exposure (corresonding to “Direct - No doors”), demonstrating the usual latent learning effect. Interestingly, when doors were activated, animals without pre-exposure (“Direct - Doors”) performed virtually identically as those with pre-exposure (“Targeted - No doors/Doors”).

In our simulations we considered the four cases studied by Daub, plus two more cases that he did not study, where doors were present during pre-exposure (“Targeted - Doors/No Doors” and “Targeted - Doors/Doors”). Analogous to time limits in the experimental study, we imposed a stricter time limit of 500 step. Our simulations reveal interesting features (Fig. 11c). First, direct and latent learning agents both failed to master the task if no doors were used (Fig. 11**c**, orange and brown curves), emphasizing the difficulty of the task under the stricter step limit. Nevertheless, the latent learning agent outperformed the direct agent during the learning phase despite reaching the goal during the pre-exposure phase only in a few trials. Second, activating doors in the learning phase vastly improved the performance of both the latent and direct learning agents (Fig. 11**c**, red and blue curves) and the learning speeds were comparable, as they were in Daub’s experiment. Third, activating doors during pre-exposure enabled the agent to master the task without doors in the learning phase, performing similarly to direct learning with doors (Fig. 11**c**, purple curve). Fourth, activating doors during both pre-exposure and learning phase simulations led to the best performance (Fig. 11**c**, green curve).

Our simulation results suggest that the activation of the doors helped the agent explore the environment more efficiently and, thus, find the goal location within the short time limit in the initial learning trials. This, in turn, helped the agents learn faster trial-by-trial than in the simulations exclusively without doors. This learning aid diminished the advantage of pre-exposure (second and third observations). Indeed, using doors during pre-exposure resulted in a representation oriented towards the following states (see Supplementary Fig. 2) that allowed our agent to learn the task without doors (third observation) within the short time limit — a task that other agents trained without doors during learning could not perform. Nevertheless, even in this setup there is room for latent learning as evident in the fourth observation.

## Discussion

The reinforcement learning (RL) agent based on deep successor representation (DSR) successfully exhibited latent learning by learning the spatial layout during pre-exposure phases. The agent encoded spatial information in two ways: a transition model representing connections between neighboring states and a forward state transition model connecting the current state directly to the goal state. The latent learning effect is modulated by the task design in the pre-exposure phase and mediated via the SR learning during pre-exposure. In contrast to other RL algorithms, such as Dyna-DQN and model-based, targeted pre-exposure improves performance more on subsequent learning tasks compared to continuous pre-exposure, because it learns a policy during pre-exposure that guides the agent toward the goal more efficiently when the reward is introduced. These results are qualitatively consistent with experimental results [5, 6].

Our DSR agent showed key differences when using one-hot encoded or image-based state representations. With one-hot encoding, the network built highly localized SF that remained tied to the training set, failing to generalize to novel inputs and resulting in scattered activations without directional bias toward the goal. In contrast, when trained on image features, which resemble what animals do, the network extracted shared structure across states, enabling generalization to unseen inputs, shown by consistent trajectories oriented toward the goal. Despite those differences, both representations consistently exhibited latent learning in all pre-exposure designs.

Our modeling results point to a key role of the exploration strategy, for which we have only very simplistic mechanisms in our model, i.e., *ε*-greedy and softmax. On the one hand, the exploration strategy helps the agent search the environment more efficiently and find the goal more quickly. On the other hand, the exploration strategy also helps build a more useful cognitive map, because the SR depends on the policy. Exploration has traditionally been considered a simple behavior compared to other spatial navigation behaviors and strategies, but more recently we have suggested that the reverse might be true and exploration might be the most complex navigation process of all [8, 24].

A way to manipulate the exploration behavior of agents/animals is to limit movement, such as by using one-way doors, which improves exploration and task performance in a spatial learning task. In these cases, doors primarily accelerate the goal finding in the early trials of the learning phase, narrowing the effect of latent learning. Our findings are consistent with Daub’s behavioral study [4], particularly the similar time scores observed between direct and latent learning animals that learned the task with doors.

Ideally, a cognitive map for an environment should encode spatial information as richly as possible, including route connections, locations of landmarks, distance and direction relations, etc. Even non-spatial attributes such as emotional associations can be stored as well [25]. However, given the limited computational resources in the brain, a cognitive map should balance its usefulness and encoding/decoding efficiency. The SR algorithm offers a trade-off between the two by encoding the temporal and spatial distance between each pair of states in the environment. It enables the animal to quickly generate a route to a given goal location when a reward is delivered.

The cognitive map has been suggested to reside in the hippocampus almost since the discovery of place cells in the hippocampus [26]. Recently, the SR has been suggested as an alternative view to place cell coding in the hippocampus [13]. Using the SR as a cognitive map, hence, closes the loop involving hippocampus, place cells, SR, and the cognitive map. However, the spatial information in SR is represented rather implicitly in such a way that the transition dynamics of the external world are not captured, i.e. the next state from a state-action pair is not stored, unlike a graph representation. Therefore, flexible planning such as generating and comparing multiple routes are not straightforward to implement by using SR. For instance, it is hard to explain the replay of sequences of place cells [23, 27], if states in SR are not explicitly connected. Nevertheless, SR offers a useful and efficient representation of the external environment, and its biological basis needs further investigation.

Our modeling study makes multiple predictions for the neural mechanisms underlying latent learning. First, neural representations of space are learned in the absence of a navigation task. Surprisingly, there is little experimental evidence on this because in traditional place cell recording studies animals are rewarded for running back and forth on a linear track or chase food pallets in an open arena. Second, the nature of the spatial representation depends on the task-design of the pre-exposure phase. Third, the spatial representation of the environment at the end of pre-exposure correlates with the degree of latent learning later on. This correlation is independent of the exploration strategy.

In summary, this work contributes to a deeper understanding of the mechanisms of latent learning and its underlying neural mechanisms. It potentially explains how such learning shapes cognitive maps and influences their efficacy in spatial tasks.

## Acknowledgements

We thank Dr. Nicolas Diekmann for his valuable contributions to the CoBeL-RL framework used in this work.

## 1 Supplementary Material

**Supplementary Figure 1.**
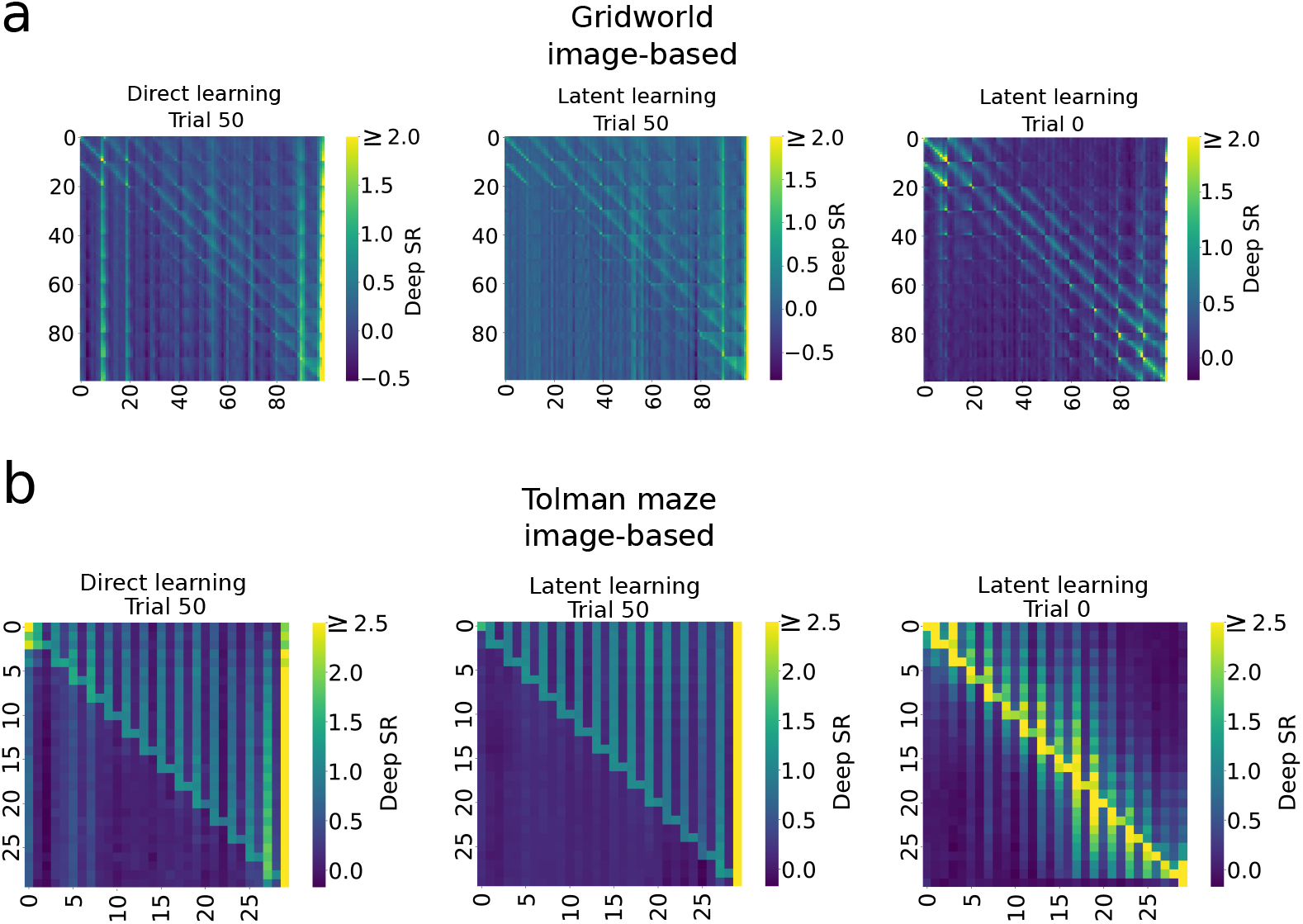
Deep SR reveal the effect of latent learning in image-based representation. SF for all states are shown for the **a**) gridworld for the right action and **b**) Tolman maze for the up action, averaged over 30 simulations. Each row of panels corresponds to the successor transitions of a particular state to every state in the environment. We computed the contribution of each state’s feature to the spatial transition using Eq. 12. Trials 0 and 50 correspond to before and after the learning phase, respectively. The SF shows that in both mazes the latent learning agent has learned the local connectivity of the maze during the pre-exposure phase before the learning phase commences. At the end of the learning phase, both agents exhibit similar transition patterns that reflect a movement from the start to the goal location in both mazes.

**Supplementary Figure 2.**
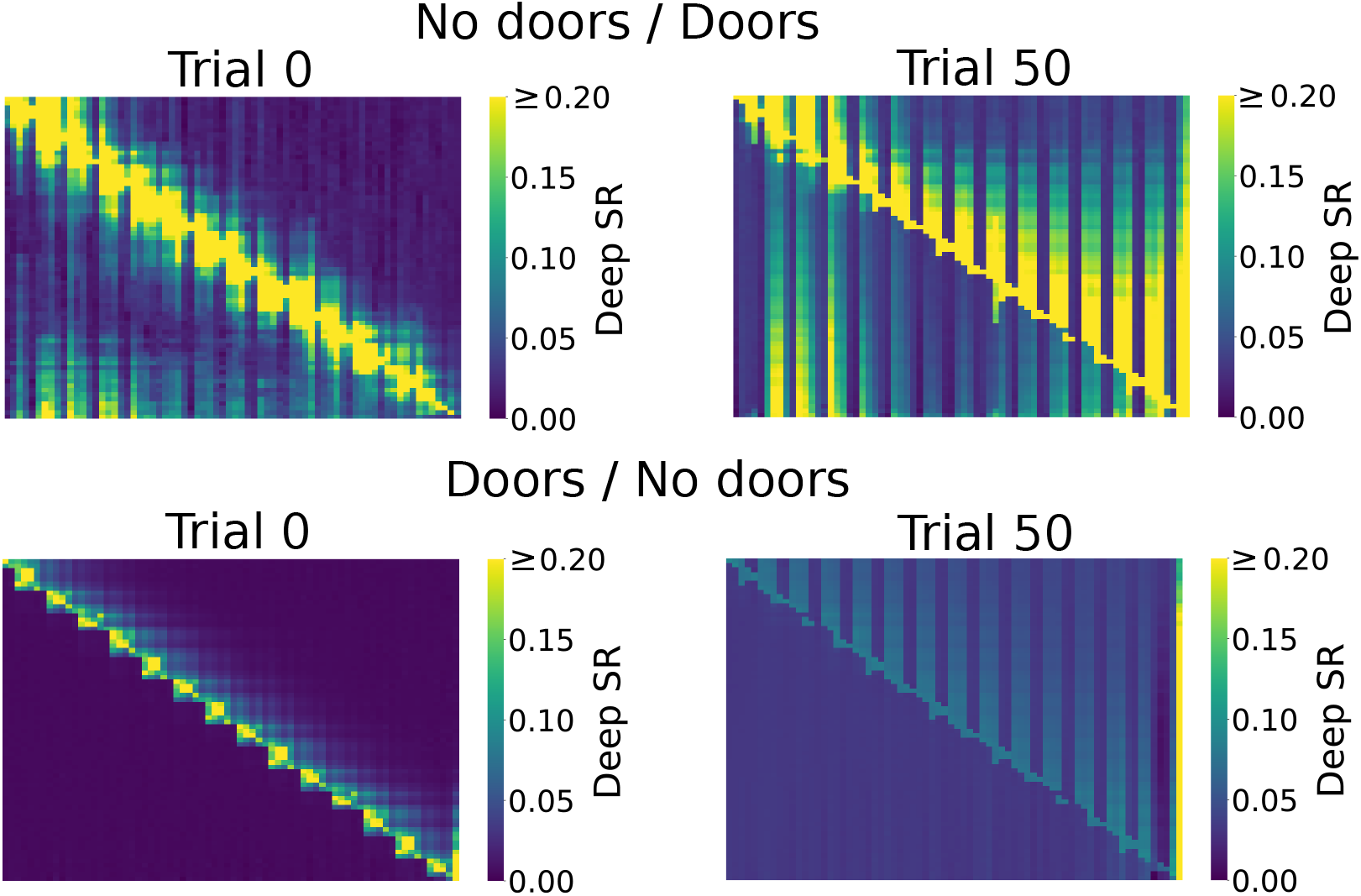
Doors facilitate the SR acquisition. Top: Deep SR of agents under No doors/Door exploration strategy. Bottom: Deep SR of agents under Doors/No doors exploration strategy. Trials 0 shows the path transition differences before the learning phase. The strategy with no doors during the learning phase shows a more concise path transition compared with the one that adopts doors.

